# Deterministic Cell Pairing with Simultaneous Microfluidic Merging and Sorting of Droplets

**DOI:** 10.1101/2025.06.27.662036

**Authors:** Kevin M. Joslin, Sophia Dateshidze, Seung Won Shin, Adam R. Abate, Iain C. Clark

## Abstract

Cell–cell interactions drive immune activation, tissue repair, and stem cell fate, yet there are few methods that can create large numbers of pre-defined cell pairs to study cell crosstalk. Droplet microfluidics allows high-throughput compartmentalization of multiple cells, but random loading results in <1% of droplets containing the desired cell combinations. Here, we present a microfluidic device capable of deterministically isolating specific cell pairs using droplet merging and sorting (‘merge-sorting’). The system detects target combinations using fluorescence and triggers simultaneous electrocoalescence and dielectrophoretic sorting. Using fluorescent dye– loaded droplets, we achieved 98.6% purity of merged and sorted droplets. In experiments using cells stained with three distinct dyes, >90% of desired cell pairs were recovered—compared to fewer than 1% when using random Poisson loading. To demonstrate the utility of this platform for extended co-culture studies, we merged cells in an alginate solution with calcium chloride droplets, producing monodisperse alginate hydrogels in which 92% of the beads contained target cell pairs that maintained viability over 18 hours. Compared to selective merger, this approach physically isolates desired droplets, eliminating unmerged contaminants and enabling cleaner downstream workflows. The device allows off-chip pre-incubation of droplets before pairing, the merger of reagents for multi-step assays, and the isolation of desired droplet pairs — capabilities not jointly accessible with existing approaches. In summary, merge-sort is a flexible platform to enrich specific combinations of droplets, cells, or particles for high-throughput studies of cell crosstalk.

## Introduction

Cell interactions play diverse and essential roles in biology, including in training the immune system to recognize pathogens^1–3^, initiating transcriptional programs that control cell differentiation^4,5^, and maintaining tissue homeostasis by inducing proliferation or apoptosis^6,7^. The diversity of cell types leads to a vast number of possible interactions, requiring systems that are both high-throughput and single-cell resolved^8–10^. To address this need, in vitro systems for compartmentalizing and assaying cell combinations have been developed. Microwells, valves, and traps have been used to create pico- and nano-liter chambers for cell culture and analysis^11–19^. These platforms have enabled co-culture and differentiation studies; for instance, embryonic stem cells have been cultured in poly(ethylene glycol) microwells, and tumor–endothelial and neuronal interactions have been studied in microvalve-controlled chambers^18,19^. However, culture in wells, even when miniaturized through microfabrication, remains relatively low-throughput because cell pairs are retained on-chip and the number of compartments is limited by the physical dimensions of the device.

Droplet microfluidics is an alternative to chip-based co-culture systems that can be used to create picoliter water-in-oil chambers for cell culture. Droplet microfluidics addresses the throughput limitations of arrayed systems by pairing cells in droplets that can be produced at kHz frequencies and collected in large numbers for bulk incubation off-chip. Mammalian cells can maintain viability in droplets over short timeframes, and the pairing of different cell populations within droplets has recently been utilized to study immune activation, cytotoxicity, and inflammation^20–25^. A critical limitation of cell-cell interaction screens in droplets is the loading of cells, which is governed by Poisson statistics and results in many drops that contain zero or one cell. This limits the number of cell pairs that can be generated, reducing the throughput of screens.

For example, in a cell pairing experiment in which two cells are each loaded at an expected frequency of 10%, only approximately 1% of droplets will contain a pair of interest. One way to address this issue is to immediately droplet-sort cell pairs on-chip using pneumatic, acoustic, magnetic, or dielectrophoretic (DEP) forces^26–34^. However, this approach is not compatible with all experimental designs. Studies involving secreted factors or requiring one cell to be treated or activated often necessitate pre-incubation in droplets prior to pairing^35–39^. Other workflows require the timed addition of reagents—for example, introducing assay components that are incompatible with cell growth at earlier stages. A microfluidic platform that enables deterministic pairing of pre-incubated droplets would provide greater flexibility and throughput for studying cell–cell interactions.

Merger of paired droplets is a powerful method for combining reagents in multi-step workflows that use cells, enzymes, or chemicals^40–46^. One significant advantage of this approach is that it keeps droplets isolated until the moment of a merger, thereby precisely controlling when their contents are mixed and reactions occur. For instance, fast-reacting hydrogel precursors like calcium ions and sodium alginate immediately gel and clog if co-flowed in microfluidic channels without fluidic shielding, but can be combined by droplet merger^47–49^. Cells cultured in droplets can be merged with assay reagents incompatible with normal cell growth^20,50–53^. Cells lysed with proteinases and heat-inactivated in droplets can be merged with PCR reagents for subsequent single-cell DNA amplification^54–56^. However, despite the advantages of droplet merger, it does not overcome the limitations of random loading of cells, beads, or molecules. Even if droplet merger is triggered based on the presence of cells^57^, unmerged drops will remain after collection and need to be processed in the subsequent steps^25^.

Here, we present a microfluidic device that merges and sorts selected droplets in a single step, a process we refer to as merge-sorting. This platform allows droplet pre-incubation, the subsequent addition of incompatible assays or chemicals, and the isolation of cell pairs of interest. Our device leverages a laser-PMT detection system^58^ and dielectrophoresis to merge and sort cell-containing droplets based on their fluorescence, overcoming the limitations of Poisson loading and creating populations that are highly enriched for the desired cell combinations. We demonstrate the utility of this approach by merge-sorting cell pairs to create cell-laden alginate beads. These beads, formed via droplet merger of alginate and calcium-containing droplets, can be recovered from oil and incubated in media, supporting cell viability for at least 18 hours and allowing downstream functional or imaging assays. Unlike selective merger approaches, which leave unmerged alginate and calcium droplets in the emulsion, merge-sorting removes background droplets, preventing nonspecific gelation during bead recovery. In summary, we present a microfluidic device that simultaneously merges and sorts droplets to produce and isolate defined cell pairs in real time. Merge-sorting is compatible with off-chip incubation and eliminates unmerged contaminants that can interfere with downstream workflows. We anticipate this device will be a valuable addition to multi-step droplet microfluidic pipelines, particularly those for studying cell–cell interactions.

## Results and Discussion

To simultaneously merge and sort target drops, we modified our concentric sorter design (34) to include a drop maker upstream of the drop re-injector (**Fig. 1a**). The device includes a saline moat (A, B) to shield drops from coalescence and prevent premature merging, a saline electrode (C) to initiate droplet merging and sorting, a droplet generator (D,E), and a port (F) for re-injecting droplets. Spacer oil (G) is used modulate the distance between droplets as they enter the sorter, bias oil (H) is added to ensure unsorted droplets exit through the waste outlet (J), and air (I) is injected to push sorted droplets through the collection outlet (K) and into the outlet tubing (**Fig. 1a**). To operate the device, drops made using a standard co-flow microfluidic chip are re-injected and paired 1:1 with on-chip generated droplets (**Fig. 1b**). Droplet fluorescence is detected using a droplet cytometer system previously described^58^. Target droplet pairs are merged by activating the on-chip electrode with a high frequency pulse, and simultaneously sorted into the sort channel using a concentric electrode (**Fig. 1b**). When the electrode is turned off, drops are continuously paired (**Fig. 1c, top**) but do not merge and flow into the waste outlet. When the electrode is activated, gated droplet pairs are merged and sorted into the sort channel (**Fig. 1c, bottom**), resulting in the collection of homogeneous, merged droplets.

**Figure 1.**
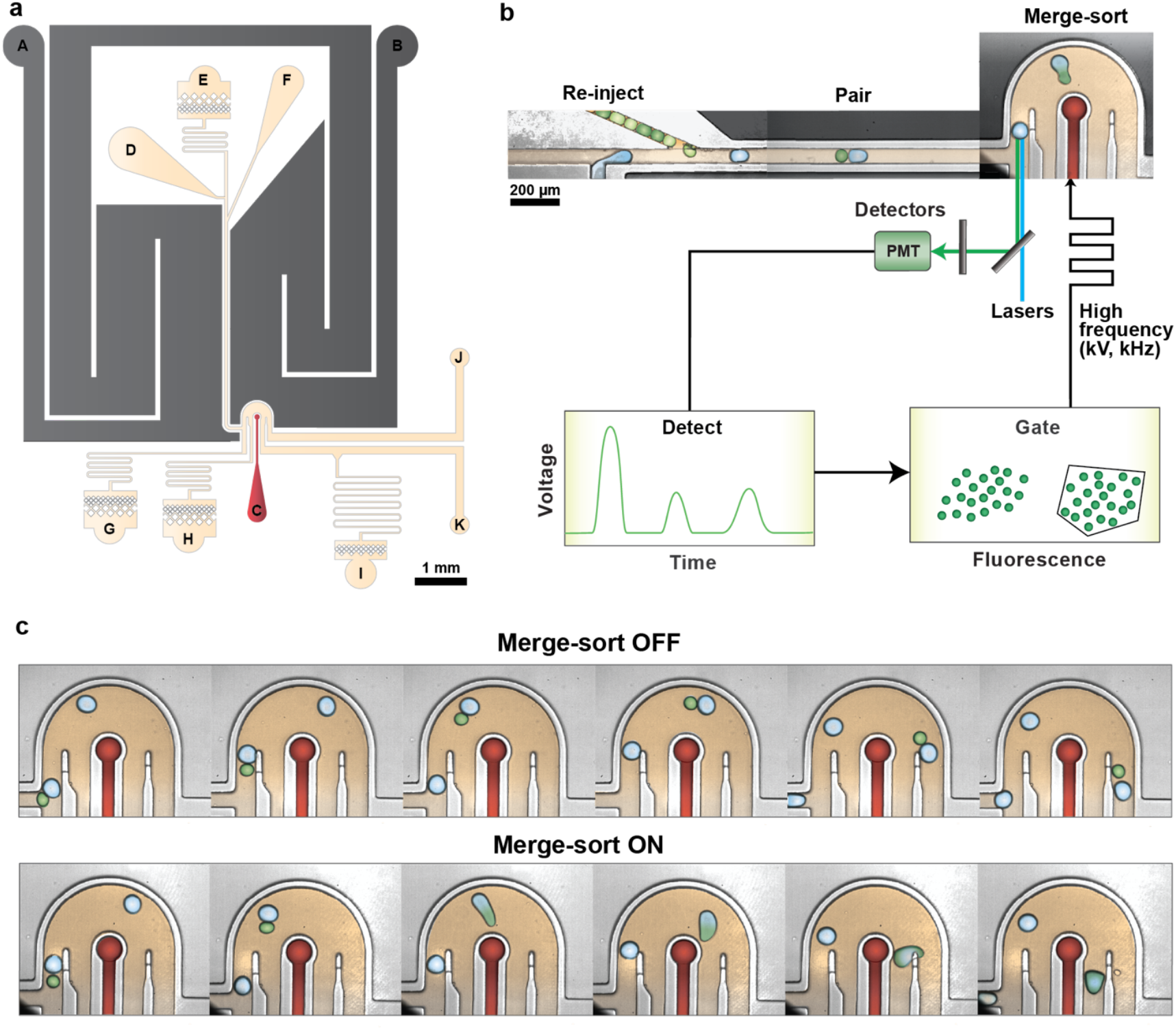
Overview of the merge-sort device design and operation: **(a)** Colorized schematic of the microfluidic merge-sort device with annotated features. A: moat inlet; B: moat outlet; C: electrode; D: cell inlet; E: surfactant oil; F: drop reinjector; G: oil spacer; H: bias oil; I: air inlet; J: waste outlet; K: collection outlet. **(b)** Colored schematic of drop pairing, merging, and sorting, and associated laser-PMT-FPGA detection system, which is used to detect droplet fluorescence and activate the saltwater electrode with a high voltage/frequency pulse. Fluorescent droplets are simultaneously merged and sorted into the collection outlet. **(c)** Time-lapse images of the device operating when the electrode is turned OFF (top) and when the electrode is triggered ON (bottom).

To test the efficiency of the device, we prepared and mixed two populations of small 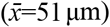 droplets containing different levels of fluorescence (FITC++, 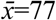; FITC-, 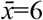 (**Fig. 2a**). We re-injected these droplets onto the device, paired them with a large, non-fluorescent droplet generated on-chip, and merge-sorted only the FITC++ population. We collected and analyzed droplets from the waste channel when the electrode was off (**Fig. 2b**), and from the sort channel when FITC++ drops were merged and sorted (**Fig. 2c**). As expected, when the merge-sort electrode was off, the collected drops consisted of unmerged small FITC++ drops, unmerged small FITC-drops, and unmerged large FITC-drops (**Fig. 2b**). When FITC++ droplets were merged and sorted, 98.6% (n=936) of droplets collected from the sort channel were FITC+, had the expected dilution of FITC fluorescence 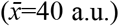, and had an increase in size 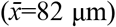 resulting from droplet merger (**Fig. 2d,e**). Together, these results demonstrate that the device can accurately merge-sort droplets.

**Figure 2.**
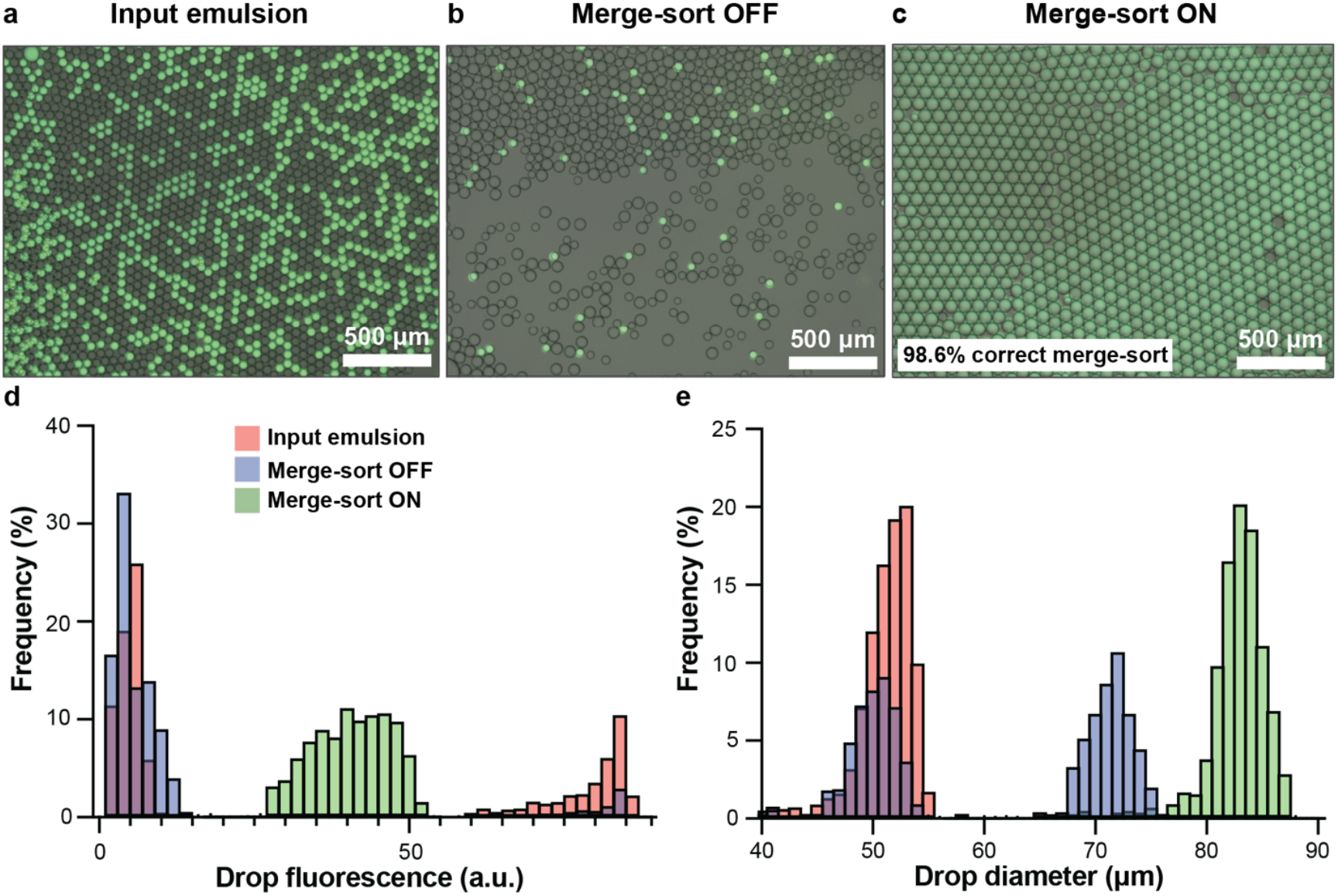
Performance of the merge-sort device using distinct populations of fluorescent droplets. **(a)** Input emulsion containing droplets with high (FITC++) and low (FITC-) fluorescence. **(b)** Collected emulsion when the electrode was OFF. **(c)** Collected emulsion when the electrode was ON. 98.6% of droplet pairs were merge-sorted correctly. **(d-e)** Histograms of the measured drop fluorescence (d) and drop diameter (e) for the input emulsion (red), merge-sort OFF collected droplets (blue), and merge-sort ON collected droplets (green).

Generating droplets containing multiple cell types results in a low percentage of the desired cell pairs, especially from within a heterogeneous sample. For example, at an average loading of 10% (λ=0.1) for each cell type, less than 1% of droplets will contain both cells, limiting the throughput of downstream analysis. Merge-sort is a powerful approach for creating cell pairs for high-throughput assays, especially cell-cell interaction screens that require large numbers of cells. To demonstrate the use of merge-sort for this application, we separately stained K-562 cells with three dyes (Cell 1, Calcein Green AM, “green”; Cell 2 CellTrace Yellow, “red”; Cell 3 CellTrace Far Red, “magenta”). We encapsulated Cell 2 and Cell 3 in drops (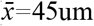um diameter) at an average loading of ~10%. Next, we re-injected and paired this emulsion with Cell 1 containing drops generated on-chip (**Fig. 3a, Video S1**), using merge sort to create the desired cell pairs (**Fig. 3b, Video S2**). This generated drops that were highly enriched for Cell 1 & Cell 2 (90% green/red, n=103) (**Fig. 3c**), or Cell 1 & Cell 3 (94% green/magenta, n=326) (**Fig. 3d**). As expected, when all droplets were merged, regardless of occupancy, there was no enrichment for specific cell combinations and the resulting emulsion contained mostly empty and single-occupied droplets (**Fig. 3e**). When the electrode was completely turned off, no merging or sorting occurred, and no Cell 1 & Cell 2 or Cell 1 & Cell 3 pairs were observed (**Fig. 3f**). These results demonstrate the ability of merge-sort to deterministically pair and isolate cell combinations at high purity (**Fig. 3g**). In addition, as we demonstrate, it is possible to successively create different cell combinations from the same sample.

**Figure 3.**
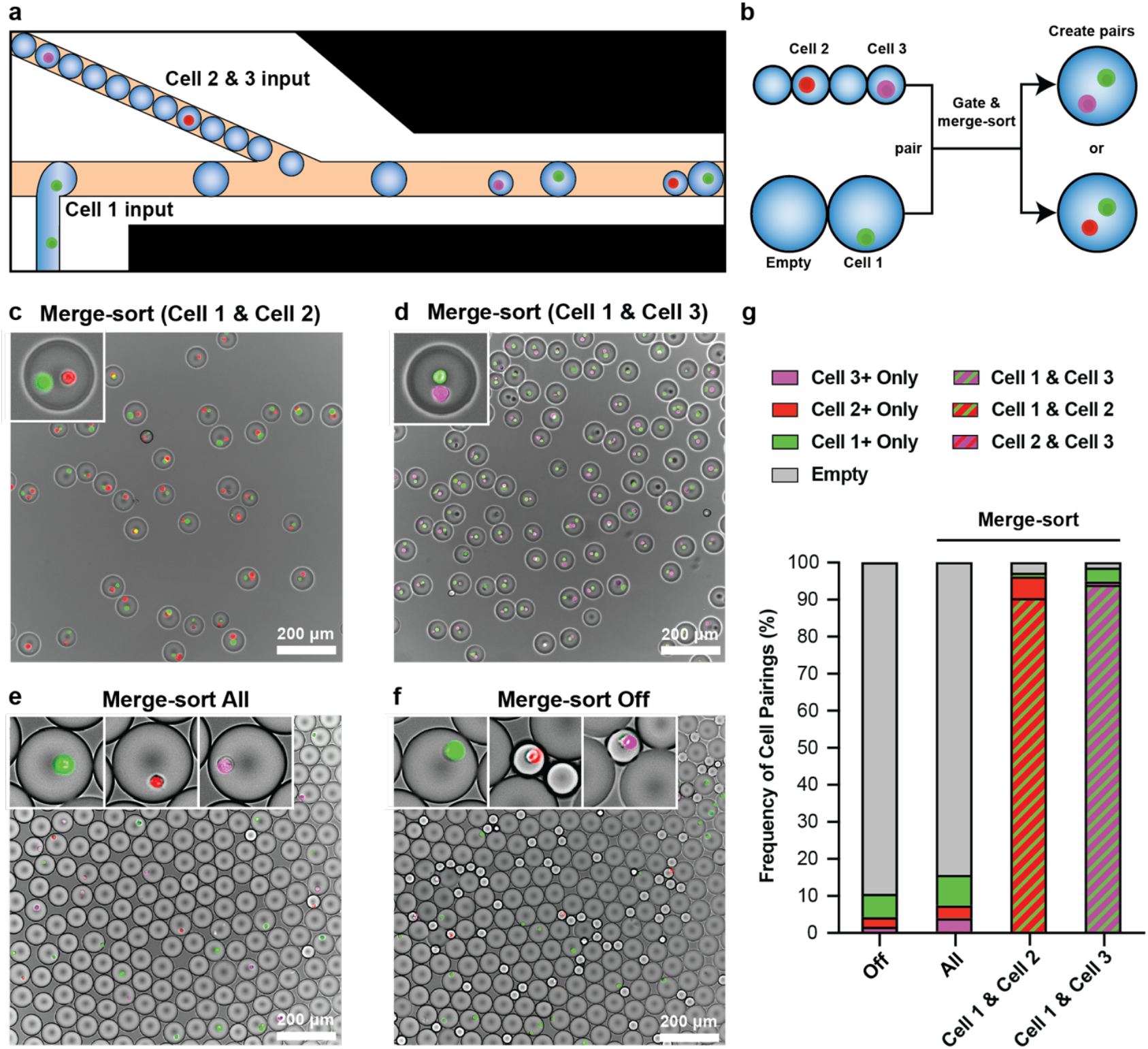
Enrichment of specific cell pairs in microfluidic droplets. **(a)** Representative illustration of re-injected droplets containing K-562 cells stained with CellTrace yellow (CTY, Cell 2, “red”) or CellTrace Far-red (CTFR, Cell 3, “magenta”). These cells are paired one-to-one with on-chip generated drops containing K-562 cells stained with Calcein Green, AM (CG, Cell 1, “green”). **(b)** Schematic depicting the pairing, selection, and generation of merge-sorted drops with Cells 1 & 2, or Cells 1 & 3. **(c-f)** Droplets collected during merge-sort selection of Cells 1&2 (upper left), merge-sort selection of Cells 1 & 3 (upper right), merge-sort selection of all droplet pairs (lower left), and when the merge-sorter is turned off (lower right). **(g)** Bar charts displaying the frequency of cell pairing when merge-sorting is turned off (Off), when all droplet pairs are merged (All), when Cells 1 & 2 are merge-sorted, or when Cells 1 & 3 are merge-sorted.

While generating cell pairs is also possible by selectively merging droplets without sorting, this has several disadvantages in screening and genomics applications. Selective merger does not remove empty drops or those with single cells, and the resulting emulsion still contains a low frequency of desired pairs. This can be especially problematic if the analysis of cell pairs includes imaging, in which 100 drops might need to be sampled to find a single cell pair. Another significant disadvantage of selective merger is that all drops, merged and unmerged, are collected in the same tube. This means that significant mixing of drop contents occurs during material recovery for downstream analysis. Merge-sort solves this problem by ensuring unmerged drops are not present during oil removal (drop breaking). This is important for recovering nucleic acids from target cells, for performing chemical reactions only in merged droplets, or, as we show here, for gelling alginate hydrogels with cell pairs.

Droplets do not allow for easy media exchange, making them poorly suited for cell–cell interaction studies requiring long-term culture. Encapsulating cell pairs in alginate hydrogels provides a biocompatible matrix that supports viability, growth, and assay compatibility over extended time periods. However, microfluidic alginate bead formation must circumvent the rapid on-chip gelation of alginate upon contact with calcium ions (Ca^2+^). Reported approaches address this by releasing chelated calcium by changing pH, using a shielding flow, or merging calcium with alginate drops^47–49,59–61^. None of these approaches avoid the generation of many empty and single-positive alginate beads in cell pairing applications (**Fig. 4b**). Selective merger can be used to create hydrogels with cell pairs, but calcium and alginate released from unmerged drops during oil removal results in large alginate aggregates that make the resulting hydrogels unusable (**Fig. 4c, Video S3**). Merge-sort removes the effect of these contaminating droplets and enables the recovery of alginate hydrogel beads containing cell pairs. To demonstrate this, we performed merge-sort with cells in alginate droplets and re-injected calcium droplets. Upon detection of a correctly paired cell combination (red+magenta), the electrode was triggered, resulting in merger of calcium with the alginate cell droplet and the sorting of target pairs (**Fig. 4d**). The alginate gelled inside the droplet and the emulsion was broken, releasing monodispersed cell-laden beads with an average diameter of 80.9±3.4 μm (n=440) (**Fig. 4e**). These alginate beads were easily washed and handled, and were highly enriched with red and magenta embedded cells; approximately 92% (n=982) of beads contained the desired cell combination, compared to only 1.6% (n=1969) of beads when all drops were merged (**Fig. 4f**). Cells captured in beads were 97.9% (n=189) viable before encapsulation, 92.8% (n=1667) viable directly post-breaking, and 92.7% (n=1813) viable after 18 hours of incubation in cell media (**Fig. 4g**).

**Figure 4.**
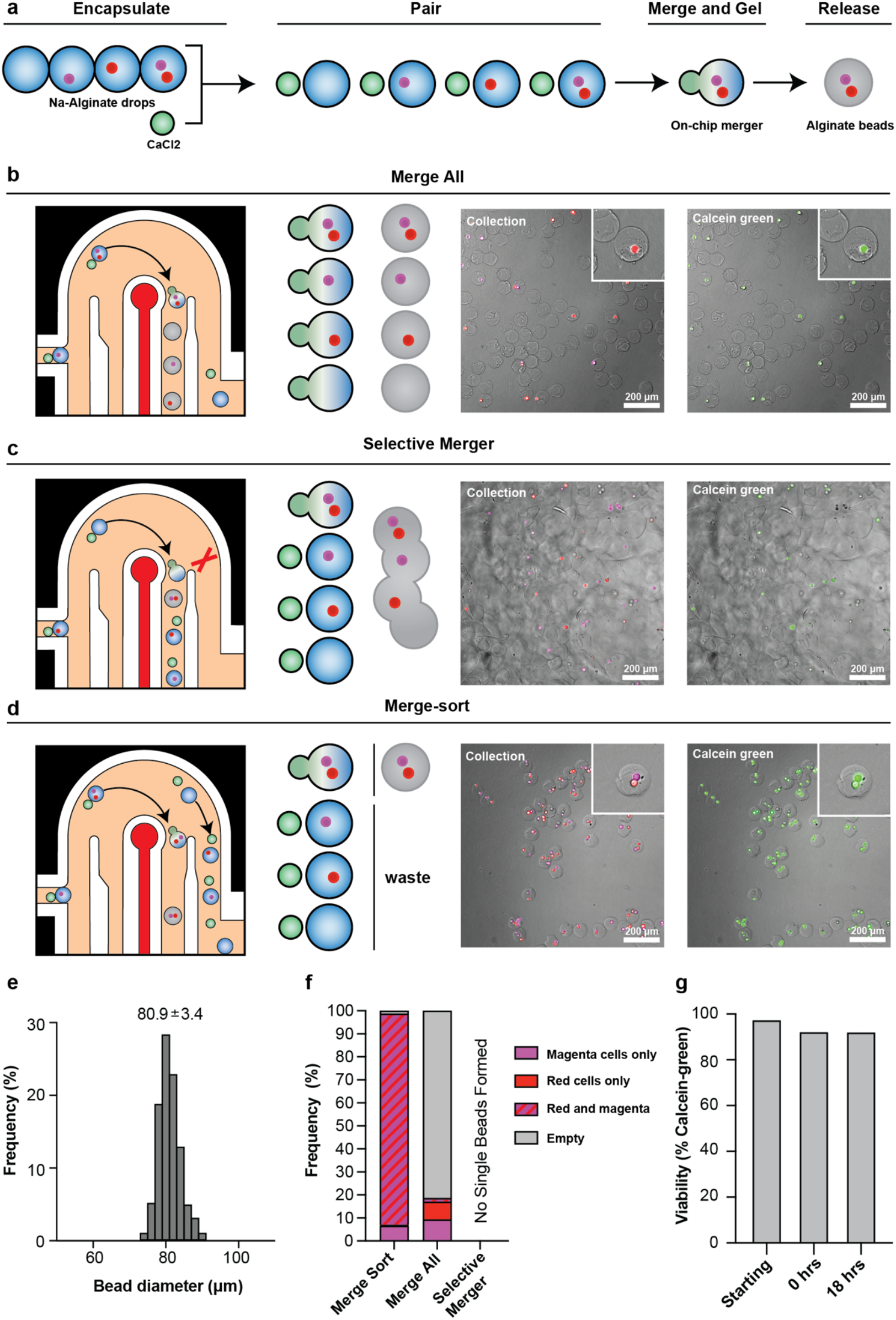
Generation of alginate beads containing desired cell pairs using merge-sort. **(a)** Schematic illustrating re-injected CaCl_2_ drops paired with on-chip generated drops containing red and magenta cells suspended in sodium alginate solution. When desired cell pairs are detected and gated, the droplets are merge-sorted, resulting in the gelation of an alginate hydrogel bead containing the selected pair. **(b-d)** Illustrations of the device operating when merging all, selectively merging, or merge-sorting (far left). Drop spacing is not to scale. Schematics demonstrating which droplet pairs are merge-sorted, merged-only, or wasted during each condition (middle left). Microscope images of cell pairing after droplet breaking for each condition (middle right). Microscope images of cell viability, measured using Calcein Green, AM stain (green), after droplet breaking for each condition (far right). **(e)** Histogram of alginate bead diameters. **(f)** Bar chart of cell encapsulation outcomes (unpaired, paired, empty) in alginate beads for each condition. **(g)** Viability of K-562 cells in alginate beads before encapsulation (starting), immediately after breaking of the emulsion (0 hrs), and after 18 hours of incubation.

Taken together, these results establish a platform for efficient and selective encapsulation of defined cell pairs in porous alginate hydrogels. Merge-sorting enables controlled mixing of calcium chloride and alginate droplets at the point of pairing, preventing premature gelation and clogging. Gelation occurs under neutral pH without the need for chelators or acidification, reducing cellular stress. The resulting hydrogels permit media exchange, maintain high cell viability after overnight incubation, and exhibit structural integrity suitable for pipette handling, making them well-suited for cell-cell interaction studies that require longer-term incubation.

## Conclusions

We present an integrated microfluidic device that simultaneously merges and sorts droplets, enabling deterministic pairing of cells, particles, or reagents. Our approach maintains isolation of droplet contents until merger, allowing off-chip preincubation and the separation of incompatible reagents before pairing. The device achieves high enrichment of defined cell pairs, far exceeding pairing frequencies from random loading, and reduces the number of droplets that must be processed in downstream assays or screens by over 100-fold. To support longer-term culture, we demonstrate the formation of alginate microspheres containing selected cell pairs with high encapsulation efficiency and viability. Unlike selective merging, merge-sorting physically separates desired droplets from unmerged ones, preventing contamination and simplifying downstream workflows. This is particularly useful for applications involving reactive chemistries or nucleic acid recovery, where the presence of background droplets can compromise results. Overall, this device provides a new droplet microfluidic tool for precise cell and reagent pairing.

## Methods

### Device fabrication

Both the co-flow drop-maker and merge-sort microfluidic devices were fabricated using standard soft lithography approaches. Specifically, photomasks were first designed in AutoCAD and custom printed (Artnet Pro). Using the photomasks and a UV light source and mask aligner, SU8 molds of the device channels were generated with SU-8 2025 or SU-8 2050 (Microchem) and 4-inch silicon wafers (University Wafer). For the co-flow drop-maker, an average layer height of 41 μm was measured by profilometry, and for the merge-sort device, average layer heights of 39 μm and 73 μm were measured for layers one and two. Poly(dimethylsiloxane) (PDMS) (Dow Sylgard) elastomer base and curing agent were mixed at a 10:1 ratio, poured on the SU8 mold, degassed, and left to cure overnight in a 65 °C oven. The cured PDMS was then cut and lifted from the mold before being punched with 0.8 mm holes and bonded to clear glass by treating with oxygen plasma and baking at 65 °C overnight. Lastly, the glass in the device was made hydrophobic with a 5 minute treatment of Aquapel (PGW Auto Glass) before immediately being flushed with air and baking at 65 °C until dry.

### Merge-sort device operation

To analyze device performance, fluorescent 50 μm drops were generated by co-flowing 1 μM acrylated FITC and 2 μM BSA-Cy5 in DPBS with Droplet Generation Oil for EvaGreen (Bio-Rad, 1864005) in a 40 μm wide standard co-flow drop-making device. The emulsion was collected into a syringe. The emulsion was re-injected onto the merge-sort device (45 μl/hr) and paired one-to-one with droplets generated on-chip. On-chip generated droplets were created by flowing 2 μM BSA-Cy5 in DPBS (200 μl/hr) and EvaGreen oil (200 μl/hr) at a T-junction. Spacer oil (1000 μl/hr) without surfactant (HFE-7500, 3M) was used to space droplet pairs, and bias oil (7000 μl/hr) was used to prevent unsorted droplets from entering the collection channel. Bubbles generated on-chip using compressed air (~3 psi) forced drops into the collection tube. A background of BSA-Cy5 is included in the droplets as a trigger for the fluorescent detection and measurement system. When a FITC- and FITC+ droplet were detected together, a 1 kV, 4 kHz, 10 cycle pulse was delivered, which simultaneously merged the droplets and pulled them into the sort channel for collection. Collected droplets were imaged on a fluorescent microscope and analyzed in ImageJ.

### Cell-cell pairing in droplets via merge-sorting

K-562 cells (ATCC) were cultured in RPMI 1640 (Gibco, 11875093) with 10% FBS and 1X penicillin-streptomycin. On the day of experiment, cells were pelleted and resuspended in 5 μM CellTrace Yellow (CTY) dye (Invitrogen, C34573), 1 μM CellTrace Far Red (CTFR) dye (Invitrogen, C34572), or 100 nM calcein green, AM (CG) (Invitrogen, C34852) in DPBS. Cells were incubated at room temperature in the dark for 20 minutes before adding 5X volume of complete media and incubating for an additional 5 minutes. Stained cells were then pelleted and resuspended in 18% Optiprep (Sigma-Aldrich, D1556) with 2 μM BSA-Cy5 in DPBS. CTY and CTFR-stained cells were encapsulated in 45 μm drops as described above and collected into the same syringe to create a mixture of droplets containing CTY or CTFR cells. This emulsion was re-injected onto the merge-sort device and paired one-to-one with CG-stained cells that were generated on-chip. CTFR+/CG+ and CTY+/CG+ cell pair droplets were detected and merged-sorted. Collected droplets were imaged on a fluorescent microscope, and cell pairs were counted in ImageJ.

### Cell-cell pairing in alginate beads via merge-sorting

Prior to the experiment, 2M calcium chloride (CaCl_2_) in water and 3% (w/v) sodium alginate (Sigma-Aldrich, W201502) in Hank’s Balanced Salt Solution (HBSS) without calcium and magnesium (Gibco) stocks were made and adjusted to pH 7.4 with sodium hydroxide. All final solutions were filtered using a 0.22 μm syringe filter. Drops 50 μm in diameter with 80 mM CaCl_2_, 1 μM FITC-acrylate, and 0.1% (v/v) poloxamer 188 (Gibco) in water were generated and collected as described above. Prior to encapsulation, K-562 cells were stained with CTY or CTFR. After staining, cells were washed once with 10 mL of 1% BSA, and 2 mM EDTA in DPBS to chelate excess divalent cations. The cells were pooled and resuspended together in 2.7% (w/v) sodium alginate, 1 μM FITC-acrylate, and 0.1% (v/v) poloxamer 188 in HBSS. Cell-containing droplets with alginate were generated on-chip, then paired one to one with reinjected CaCl_2_-containing droplets. The CaCl_2_ drops were merged-sorted with cell/alginate drops containing the desired cell pairs, resulting in gelled alginate beads. Droplets were broken and beads released by 5x v/v adding 100 mM CaCl_2_, 0.1% poloxamer 188 in water and 20% (v/v) 1H,1H,2H,2H-Perfluoro-1-octanol (PFO, Sigma-Aldrich) in HFE, followed by gentle agitation by hand. After the emulsion was broken, the supernatant containing beads was strained on top of a 40 μm cell strainer to remove excess oil and small debris. The strainer was immediately inverted, and the beads were collected into a clean tube before being strained through a 100 μm strainer to remove aggregates and large debris. The beads were then washed once in 1.8 mM CaCl_2_, 0.1% poloxamer 188, 10% FBS, 1X penicillin-streptomycin RPMI before final resuspension in the same media. Collected beads were finally stained with 100 nM CG, incubated for 15-20 minutes at room temperature, imaged on a fluorescent microscope, and analyzed in ImageJ.

## Supporting information

CAD Geometry Files

## Contributions

KMJ, ARA, and ICC conceptualized experiments. KMJ, SD, SWS, and ICC performed experiments. KMJ analyzed data. KMJ and ICC wrote the manuscript.

## Acknowledgments

ICC was supported by a grant from the NIH (DP2AI154435). KMJ was supported by the National Science Foundation Graduate Research Fellowship Program and the Hearts to Humanity Eternal (H2H8) Graduate Research Grant.

